# Electrical Propagation of Condensed and Diffuse Ions Along Actin Filaments

**DOI:** 10.1101/2021.04.03.438300

**Authors:** Christian Hunley, Marcelo Marucho

## Abstract

In this article, we elucidate the role of divalent ion condensation and high polarization of immobile water molecules in the condensed layer on the propagation of ionic calcium waves along actin filaments. We introduced a novel electrical triple layer model and used a non-linear Debye-Huckel theory with a non-linear, dissipative, electrical transmission line model to characterize the physicochemical properties of each monomer in the filament. This characterization is carried out in terms of an electric circuit model containing monomeric flow resistances and ionic capacitances in both the condensed and diffuse layers. In our studies, we characterized the biocylindrical actin filament model using a high resolution molecular structure. We considered resting and excited states of a neuron using representative mono and divalent electrolyte mixtures. Additionally, we used 0.05*V* and 0.15*V* voltage inputs to study ionic waves in voltage clamp experiments on actin filaments. Our results reveal that the physicochemical properties characterizing the condensed and diffuse layers lead to different electrical conduction mediums depending on the ionic species and the neuron state. This region specific propagation mechanism provides a more realistic avenue of delivery by way of cytoskeleton filaments for larger charged cationic species. This new direct path for transporting divalent ions might be crucial for many electrical processes that connect different compartments of the neuron to the soma.

## 1 Introduction

Innovative models and theories on the electrical activities in the cytoskeleton are becoming a valuable tool to understand information transfer inside neurons (Frieden and Gatenby, 2019; Gatenby, 2019; Frieden and Gatenby, 2020). Actin filaments and microtubules, a major component of the cytoskeleton, have gained reputation as bionanowires capable of conducting ionic waves by utilizing the biological ionic environment (Ndzana and Mohamadou, 2019). The actin-calcium relationship in neurons is crucial for many electrical processes taking place in ion channels, the dendrites, axon, terminals and the soma (Oertner and Matus, 2005; Kanwore et al., 2021; Lavoie Cardinal et al., 2005; Ganguly et al., 2015). For example, actin filaments can alter the influx of calcium at membrane voltage gated ion channels, and regulate the release of calcium deep in the nerve at the endoplasmic reticulum (Schubert and Akopian, 2004; Wang, Mattson, and Furukawa, 2002; Rosado and Sage, 2000). In addition, Tunneling nanotubes (TNTs) are a cell-to-cell connection composed of actin filaments surrounded by a lipid membrane that can transport calcium over distances up to 100 micrometers (Sartori-Rupp et al., 2019; Marzo, Gousset, and Zurzolo, 2012; Gerdes, Rustom, and Wang, 2013). Nevertheless, whether actin is a pathway for calcium to the stereocilia of the inner ear, or involved in cellular calcium signaling by storage and release, actin proteins and calcium ions have recently demonstrated to have more roles than previously understood (Tuszynski et al., 2018b; Sataric et al., 2020; Lange and Brandt, 1996). One new promising mode of ion transport inside neurons is through the cytoskeleton where actin filaments and microtubules may be a direct path for transporting divalent ions like calcium or magnesium (Sataric, Sekulic, and Sataric, 2015; Tuszynski et al., 2018a; Sataric et al., 2009; Craddock, Tuszynski, and Hameroff, 2012; Priel et al., 2008).

Experimental research under in-vitro conditions revealed that actin filaments are capable of conducting ions from one filament end to the other when submerged in an aqueous electrolyte solution (E.C. Lin, 1993; H.F. Cantiello, 1991). Theoretical models were developed to elucidate the mechanisms underlying their electrical conductivity properties using cylindrical filament models, monomeric electric circuit components, and transmission line models similar to those used to characterize action potentials propagating along axons. Unlike axons, the high surface charge of the polyelectrolyte causes an inhomogeneous arrangement of counter- and co-ions forming an electrical double layer (EDL). As a result of the intrinsic properties of the protein, and the composition of the electrolyte fluid, cytoskeleton filaments build up an ionic conductivity and capacitance layer surrounding their surfaces (Tuszynski et al., 2004; Hunley, Uribe, and Marucho, 2018). These models predict that cytoskeleton filaments are able to sustain uniform ionic waves propagating at constant velocity for distances comparable to the size of some neurons.

Recently, a multi-scale method (atomic(Å) → monomer(*nm*) → filament(*µm*)) (Fig.1) was developed to account for the charging mechanisms of residues in actin proteins, and the non-trivial contributions of the electrical double layer (EDL) to the ionic electrical conductivity along actin filaments(Fig. 2a) (Hunley, Uribe, and Marucho, 2018; Marucho, 2019). Results for in-vitro and intracellular conditions revealed changes in width, decay and velocity of propagation depending on the voltage input and conducting properties of actin filaments. However, experimental evidence on multivalent ions has demonstrated that modeling the EDL becomes inaccurate due in part to an overestimation of the relativity permittivity of the aqueous medium, a lack of consideration for immobile water molecules near the surface, and the non linear behavior of the mean electrostatic potential used in EDL theories for distances near the protein surface (Fig 2).

**Fig. 1:**
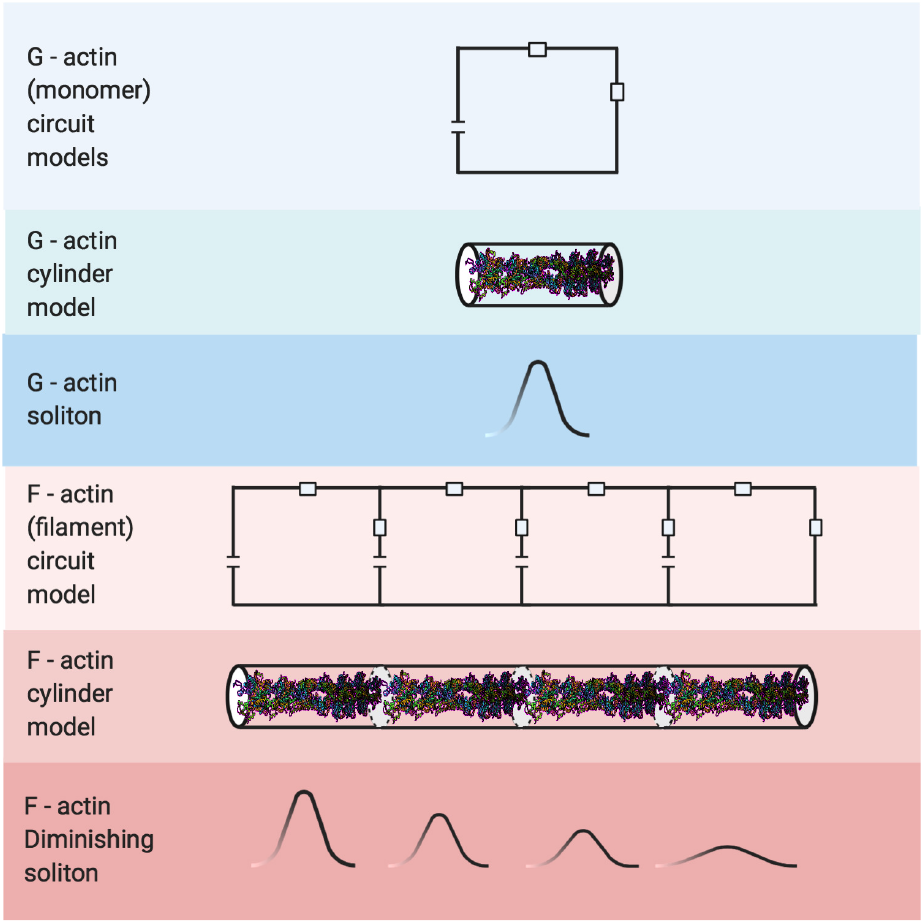
Multi-scale approach

**Fig. 2:**
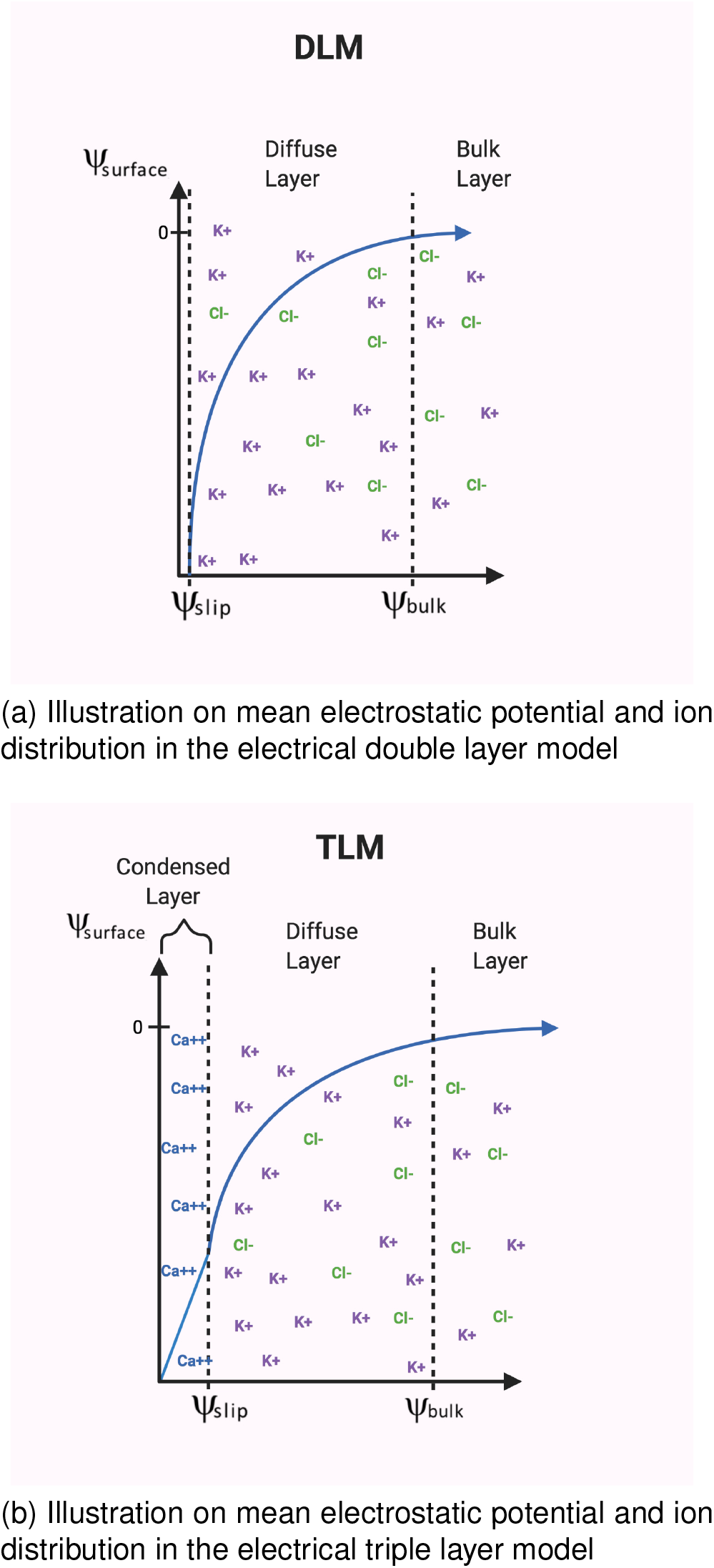
Comparison between the electrical double layer (EDL) and the electrical triple layer (ETL) models

To overcome some of these limitations, in this article, we introduce a multi-scale approach for the electrical triple layer (ETL) model to accommodate multivalent counterions in the condensed layer next to the hydration radius of the actin filament (Fig. 2ab). The ETL model considers a sigmoidal-like (step function) relative dielectric permittivity with the bulk value in the diffuse layer, and a much lower value used in the condensed layer to account for the high polarization of water molecules near the filament surface. Additionally, we consider a uniform counterion concentration in the condensed layer which is in agreement with more sophisticated theories accounting for ion-ion correlation and water crowding. The standard Boltzmann distribution for the ion concentrations and a non-linear Debye-Huckel theory are used for the diffuse layer to account for the filament charge renormalization and the condensed layer thickness (Lamm and Pack, 2010). The approach considers the immobile water molecules in the condensed layer, whereas, the water molecules in the diffuse layer move parallel to the filament following the Naiver-Stokes law. Using the ETL model we characterize the physicochemical properties of each monomer in the filament in terms of an electric circuit model containing monomeric flow resistances and ionic capacitances in both the condensed and diffuse layers. We use the resistance and capacitance combination rules to convert the ETL model into a EDL monomeric electric circuit containing equivalent resistances and capacitance. This electric unit cell is used in the non linear, dissipative, transmission line prototype recently proposed for the EDL model to investigate electrical signal propagation along F-actin immersed in mixtures of mono- and -divalent ionic aqueous solutions. Finally, we consider different external voltage inputs and calcium concentrations, which are usually present for in-vitro experiments on single neurons, to characterize the electrical impulse peak, width, and velocity of propagation, as well as, the distance and travel time of ions transversing the end-to-end filament length.

## 2 Theory

In this section, we describe the methodology used to study a triple layer model using a multi-scale approach. The notation used in this work for the mean electric potential and ionic waves is based on the articles (Lamm and Pack, 2010) and (Hunley, Uribe, and Marucho, 2018), respectively. In particular, 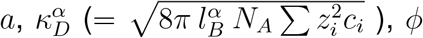, and 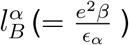 represent the filament radius, inverse Debye length, mean electrostatic potential, and Bjerrum length. Whereas *α* represents the diffuse layer (*DL*) or the condensed layer (*CL*). The variable *σ*_0_ is the bare surface charge density normalized by the fundamental electric charge, *e*, and *β* = 1*/*(*k*_*B*_*T*). The parameters *N*_*A*_, *k*_*B*_, and *T* are the Avogadro number, Boltzmann constant, and temperature in Kelvin. Further, 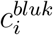 and *z*_*i*_ stand for the bulk concentration and valence of the ionic species *i*, respectively. To simplify calculations for the mean electrostatic potential we used the dimensionless variables 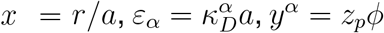, and 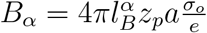. The parameter *z*_*p*_ represents the effective electrical valence for electrolytes containing both mono- and divalent ions.

### 2.1 Cylindrical Biomolecule Model

We use the cylindrical biomolecule approach for G-actins introduced in our previous article to calculate the radius and surface charge density from the molecular structure model of an actin filament (Hunley, Uribe, and Marucho, 2018).

### 2.2 Electrical Triple Layer Model

#### 2.2.1 The condensed and diffuse layers

In this work, we model a condensed layer, i.e., the region between the charged polymer and the diffuse layer, to characterize the electric conductivity along actin filaments for conditions involving divalent ions like calcium and/or magnesium. We characterize the relative dielectric permittivity in the electrical triple layer model using two values. We use *ϵ*_*CL*_ = 5 in the condensed layer to account for the strong polarization of the electric dipole of the immobile water molecules surrounding the filament (Warshavsky and Marucho, 2016). Whereas, in the diffuse layer, we use *ϵ*_*DL*_ = 78.358 which corresponds to the bulk electrolyte solution. We represent such a dependence of the relative dielectric constant on the separation distance using the following step function

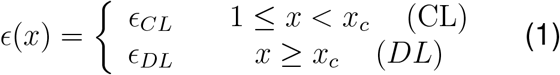

where *x*_*c*_ − 1 stands for the condensed layer thickness. This model is in agreement with the polar solvent density functional theories that predict a sigmoidal-like distance dependent relative dielectric constant *ϵ* (*x*) (Warshavsky and Marucho, (2016)). In these theories, distances with larger separation from the surface have a bulk solvent value, while a lower value is more accurate near the surface of the charged wall. It is worth noting that *ϵ*_*DL*_ = 78.358 is the value usually used in conventional approaches that only consider the diffuse layer as the propagating media for ionic waves.

Additionally, the competition between the short-range ion-ion repulsion and water crowding forces overcome the long-range electrostatic interactions near the surface of charged objects in aqueous electrolytes (Hunley and Marucho, 2017). The result of this competition is a break down in the Boltzmann statistics (Ren et al., 2012), and leads to a rather uniform counterion concentrations in this very thin condensed layer. Whereas, the Poisson-Boltzmann equation is still accurate to describe the mean electric potential and ion distributions in the diffuse layer where the electric potential force dominates, and the monovalent counterion concentration falls off with a nonlinear decreasing monotonic behavior (Fig. 2). Thus, in this work, we use the following Poisson equation to estimate the mean electrostatic potential and ion distributions in both the condensed and diffuse layers

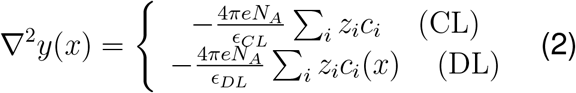

where the CL ranges from 1 ≤ *x* ≤ *x*_*c*_, and the DL from 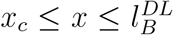. In equation (2), *c*_*i*_ is the concentration of the counterion species in the CL, and 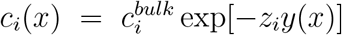 is the Boltzmann concentration distribution of all ion species in the DL. We solve the Poisson equation (2), along with the following standard boundary conditions at the interface of the two layers: ∇*y*(*a* = 1) = −*B*_*SL*_, *y*^*DL*^(∞) = 0.

Additionally, we use the non-linear Debye-Huckel approximation introduced by Lamm and Pack to account for the ion condensation (Lamm and Pack, 2010). The approach generalizes the linear approximation, providing an analytic, yet accurate expression for the mean electrostatic potential in terms of an effective surface charge density, and has been shown to be in qualitative agreement with Manning’s condensation theory (Lamm and Pack, 2010; Manning, 2007; Manning, 2011). We also consider a symmetric *z*_*p*_ : *z*_*p*_ electrolyte solution with effective ionic valence 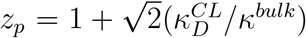 to approximate the mean electric potential generated by an asymmetric (1 : 1 − 2 : 1 or 1 : 1 − 2 : 2) electrolyte mixture of mono- and divalent ions.

These models, theories, and approximations on the ETL yield the following expression for the dimensionless mean electric potential in the diffuse layer

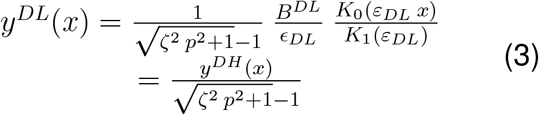

Interestingly, the mean electric potential in the diffuse layer of the TLM, *y*^*DL*^(*x*), differ from the electric potential, *y*^*DH*^ (*x*), used in our previous work by a factor 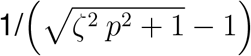. This implies that the ion condensation in the TLM generates an effective surface charge density *σ*, which is related with the bare surface charge density *σ*_*o*_ in our previous model as follows 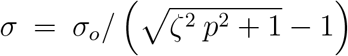. This expression is a generalization of the result predicted by the two states of Manning’s condensation theory. The parameter *ζ* is calculated numerically from the equation 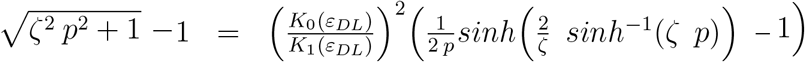, where the variable *p* is related to the dimensionless mobility and relative dielectric permittivity of the diffuse layer as 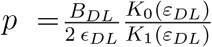.

The corresponding equation for the mean electrostatic potential *y*^*SL*^(*x*) in the condensed layer is given by

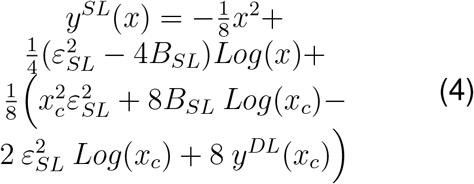

#### 2.2.2 Triple Layer Structure

The bjerrum length is usually used to estimate the electrical double layer thickness since it represents the radial distance from the protein surface to the location where the electrostatic and thermal energy become equivalent. In the triple layer model, we split this region into a condensed layer (1 ≤ *x* ≤ *x*_*c*_) which contains the multivalent counterions, and a diffuse layer (*x*_*c*_ ≤ *x* ≤ *l*_*B*_*/a*) containing monovalent counter- and co-ions.

The condensed layer thickness (*x*_*c*_ − 1) is obtained from the electro-neutrality condition using the following equation

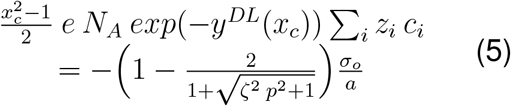

### 2.3 Electric Circuit Elements in Both the Condensed and Diffuse Layers

In this section, we use the fluid transport laws and the analytic expressions obtained for the mean electric potential to calculate the total transversal and longitudinal currents for both the condensed and diffuse layers of the triple layer model.

#### 2.3.1 Longitudinal (axial) Current and Resistance

The expression for the longitudinal current density *i*_*l*_(*x*) = *k*(*x*)*E*_*z*_ + *v*_*z*_(*x*)*ρ*_*e*_(*x*) (electro-osmotic plus natural convection) depends on the triple layer structure since it considers immobile and mobile water molecules in the condensed and diffuse layers, respectively. Thus, we neglect the convection term 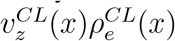 in the condensed layer. The explicit expressions for *v*_*z*_(*x*) and *ρ*_*e*_(*x*) in the 9diffuse layer are 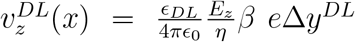 and 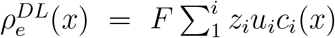. Additionally, 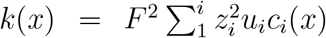 represents the effective electrolyte conductivity where *c*_*i*_ is the ion concentrations contained in their respective layer. As a result, we can write 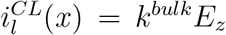 and 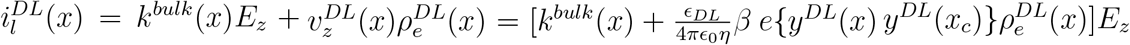.

Additionally, the upper and lower limits of integration used to calculate the total longitudinal currents *I*_*l*_ = 2*πa*^2^ ∫ *i*_*l*_(*x*)*xdx* depend on the triple layer structure. Accordingly, we have 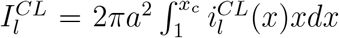 and 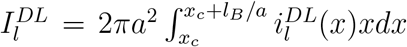. Further, 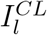 and 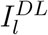 circulate in parallel with each other since Δ*V*_*DL*_ = Δ*V*_*CL*_ = |Δ*V* |= *E*_*z*_*ℓ* along the axial direction. After performing lengthy integral calculations, we get Ohm’s law 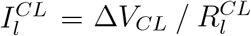 and 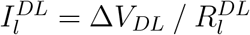 where

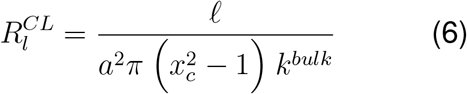

and

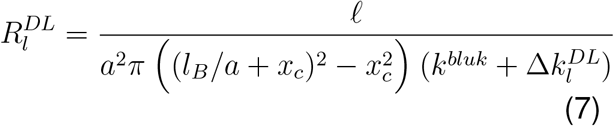

In the latter equation 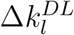 represents the non trivial contributions to the bulk electrical conductivity coming from the diffuse layer. Its explicit expression is included in the appendix.

#### 2.3.2 Transversal (radial) Current and Resistance

The expression for the transversal current density 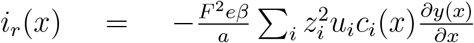 depends on the triple layer structure since we only consider counterions in the condensed layer, whereas all ion species are present in the diffuse layer. As a result, we can write 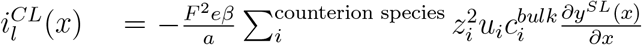 and 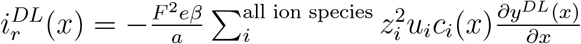.

Following the total longitudinal current calculations, the expressions for the total transversal currents 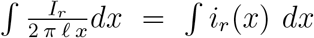 in the condensed and diffuse layers are given by 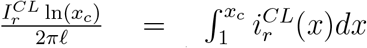 and 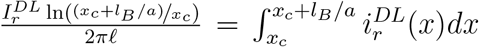, respectively. Additionally, 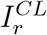 and 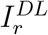 circulate in series since Δ*V*_*DL*_ = *y*^*DL*^ (*x*_*c*_ + *l*_*B*_*/a*) − *y*^*DL*^(*x*_*c*_), Δ*V*_*CL*_ = *y*^*SL*^(*x*_*c*_) − *y*^*SL*^(1), and the voltage across the electrical triple layer (ETL) is Δ*V*_*ET L*_ = Δ*V*_*CL*_ + Δ*V*_*DL*_ in the radial direction. After performing the integral calculations we get the Ohm’s law 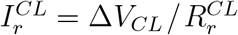 and 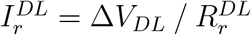 where

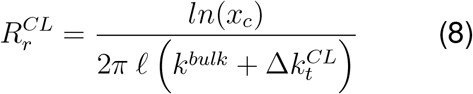

and

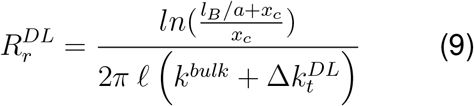

In the later expressions, 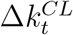 and 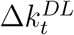 represent the non trivial contributions coming from the condensed and diffuse layers, respectively. The corresponding expressions read

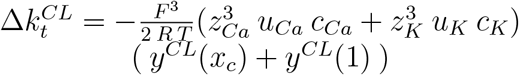

and

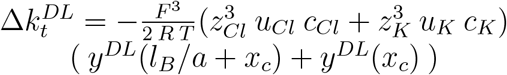

#### 2.3.3 Capacitance

In the EDL model, the charge accumulated in the capacitor has a nonlinear behavior on the applied voltage since the ion distributions in the diffuse layer satisfy the Boltzmann statistics. This remains true for the diffuse layer in the ETL model. Thus, we have

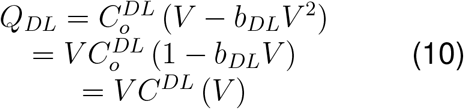

where *V* = *eβy*^*DL*^(*x*_*c*_) is the surface electric potential, 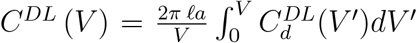 is the total capacitance, and 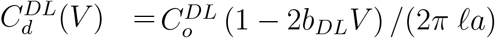 stands for the surface differential capacitance in the diffuse layer. Following the approach for the EDL, we calculate the parameters *b*_*DL*_ and 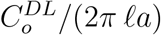 using CSDFT (Marucho, 2019).

On the other hand, the counterion concentrations are uniform in the condensed layer. Additionally, *y*^*CL*^(*x*) is a non linear function on the surface charge density due to the ion condensation. In this case, we calculate the differential capacitance 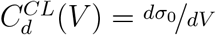 from the analytic expression obtained for the surface electric potential *V* = *eβy*^*CL*^(1) = *f* (*σ*_0_) (eqn 4). We obtain the inverse series expansion for *σ*_0_(*V*) = *f* ^*−*1^(*V*) = *a*_0_ + *a*_1_ *V* + *a*_2_ *V* ^2^ + *a*_3_ *V* ^3^ + (*V* ^4^) using the commercial software Mathematica 12.0.

As a result, the differential capacitance in the condensed layer is given by 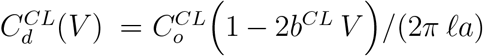 and the charge accumulated in the capacitor reads

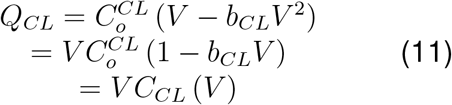

where the nonlinear parameter comes from *b*^*CL*^ = −*a*_2_*/a*_1_, and the capacitance coefficient is 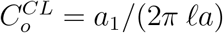.

According to the triple layer structure, the capacitors in the condensed and diffuse layers are in series since Δ*V*_*ET L*_ = Δ*V*_*CL*_+Δ*V*_*DL*_ and *Q*_*CL*_ = *Q*_*DL*_ along the radial direction. Explicit expressions for the parameters *a*_0_, *a*_1_, *a*_2_, and *a*_3_ are provided in the appendix.

### 2.4 Equivalent Electric Circuit Components and Transmission Line Model

We use the previous results to build up a monomeric electric circuit model for the TLM consisting of two resistances in parallel along the longitudinal directions, two resistances in series, and two capacitors in series along the radial direction (Fig. 3a). According to this, we apply the resistance and capacitance combination rules to convert the ETL monomeric electric circuit into an EDL monomeric electric circuit containing equivalent capacitance and resistances (Fig. 3b). The equivalent transversal and longitudinal resistances become

**Fig. 3:**
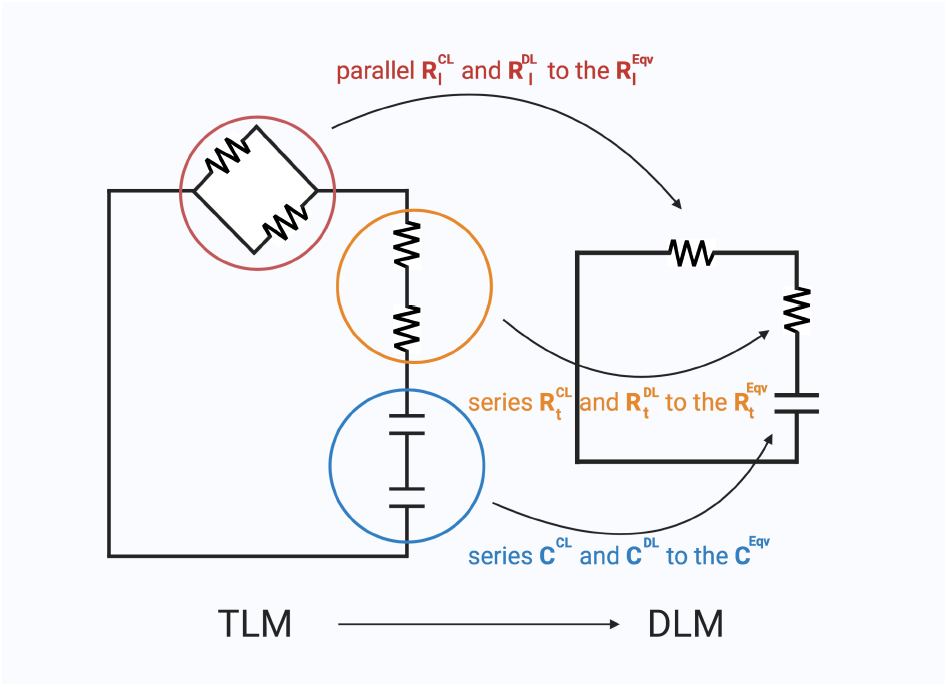
Equivalence between the Triple layer model (TLM) and the double layer model (DLM) electric circuit using the parallel and series circuit components rules.

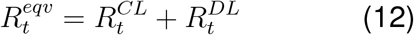

and

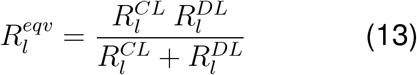

respectively. The voltage dependent equivalent capacitance is given by

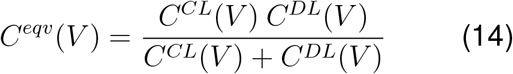

Using the expressions (10) and (11) for the capacitors *C*^*DL*^(*V*) and *C*^*CL*^(*V*) and the their corresponding Taylor expansion in powers of the voltage *V*, eq. (14) can be written as

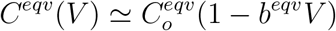

where the equivalent parameter “b” is given by

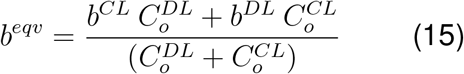

and the effective bare capacitance is

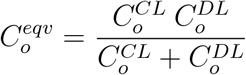

Using this equivalent electric circuit model as the unit cell in the non linear, dissipative, transmission line prototype model proposed for the EDL model, we have the following approximate analytic solution for the soliton (Hunley, Uribe, and Marucho, 2018)

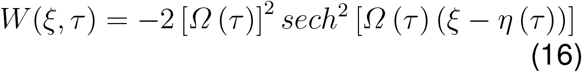

with

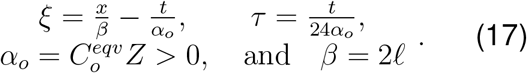

*Z*^*−*1*/*2^*U* (*x, t*) = *I*(*x, t*), and 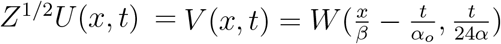

In these equations, *ℓ* is the monomer diameter, 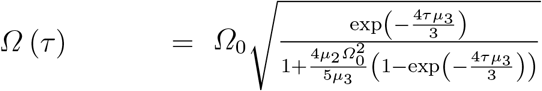 represents the time-dependent amplitude, and 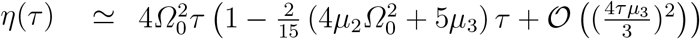. Additionally, 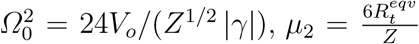 and 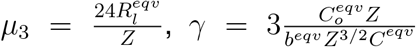, and *V*_*o*_ represents the input voltage. The expression for the impedance *Z* is provided in the appendix.

Using the previous expression for the soliton we calculate the vanishing travel time *t*_*max*_, the maximum travel distance *x*_*max*_, and the average velocity *v*_*avg*_. *t*_*max*_ is the solution of the implicit equation 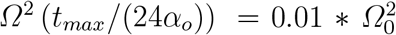. It represents the time at which the soliton amplitude attenuates to 1% of its initial amplitude, Where as *x*_*max*_ = |(1 − *t*_*max*_*/α*_*o*_)*β*| and 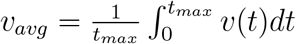 where

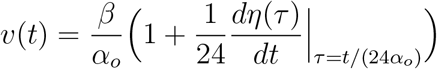

and 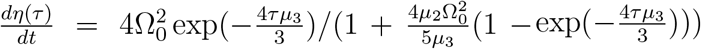.

As a unique future the TLM reduces to the DLM in the case that the thickness of the condensed layer, (*x*_*c*_ − 1), goes to zero and condensation at the surface is no longer predicted. The longitudinal resistance in the condensed layer goes to infinity acting as an electric circuit breaker, and thus, the equivalent longitudinal resistance becomes the longitudinal diffuse resistance obtained in our previous work. Similarly, the capacitor in the condensed layer goes to infinity as the separation distances between the charged layers (plates) goes to zero, and therefore, the equivalent capacitor equals the capacitor in the double layer obtained in our previous work. Whereas, the transversal resistance in the condensed layer goes to zero, and consequently, the equivalent transversal resistance equals the transversal diffuse resistance obtained in our previous work.

## 3 Results

In section 3.1, we characterize the electric circuit elements (resistances, conductivities, and capacitances) of the unit cell representing the electrical properties of the triple layer electrolyte surrounding a G-actin monomer. We also describe the triple layer properties including the thickness and fraction of the condensed ions in the stagnant layer. These results are then used in section 3.2 to characterize the propagation of ionic currents (or voltages) in form of solitons (velocity, attenuation, traveling distance, and vanishing time) along a dissipative non linear transmission line electric circuit representing the actin filament. To model the biological environment, we used the mixture of salts *KCl*+*CaCl*_2_ with a concentration range that is consistent with intracellular physiological conditions in a neuron. The resting state of a neuron is modeled with 0.1 *M K*^+^ and 50 *nM Ca*^2+^, whereas the excited state is modeled with 0.1 *M K*^+^ and 1 *mM Ca*^2+^. Additionally, we consider voltage inputs of 0.05*V* and 0.15*V* to investigate ionic waves in voltage clamp experiments on actin filaments. We also present, when appropriate, comparisons between the triple and double layer electrolyte model predictions to reveal the role of the condensed ion layer in the propagation of solitons along actin filaments.

Finally, in section 3.3, we characterize solitons in neurons going from the resting to excited states.

### 3.1 Monomeric Characterization of G-Actin

The width of the condensed layer and fraction of the condensed ions at different calcium bulk concentrations are presented in Figure 4 and Table 1a, respectively. Whereas, Table 1b shows the corresponding values predicted by Manning’s theory (Manning, 2007; Manning, 2011).

**Table 1:** Impact of bulk calcium concentration on the percent of ions condensed in the condensed (CL) and (DL) double layers. (a) and (b) represent the results predicted by this work and Manning’s theory, respectively.

| $[Ca^{2+}]$ | SL | DL | $[Ca^{2+}]$ | DL |
| --- | --- | --- | --- | --- |
| 50 nM | 7.65 | 92.35 | 50 nM | 61.72 |
| 50 $\mu M$ | 8.22 | 91.78 | 50 $\mu M$ | 63.31 |
| 1 mM | 10.18 | 89.82 | 1 mM | 68.06 |

**Fig. 4:**
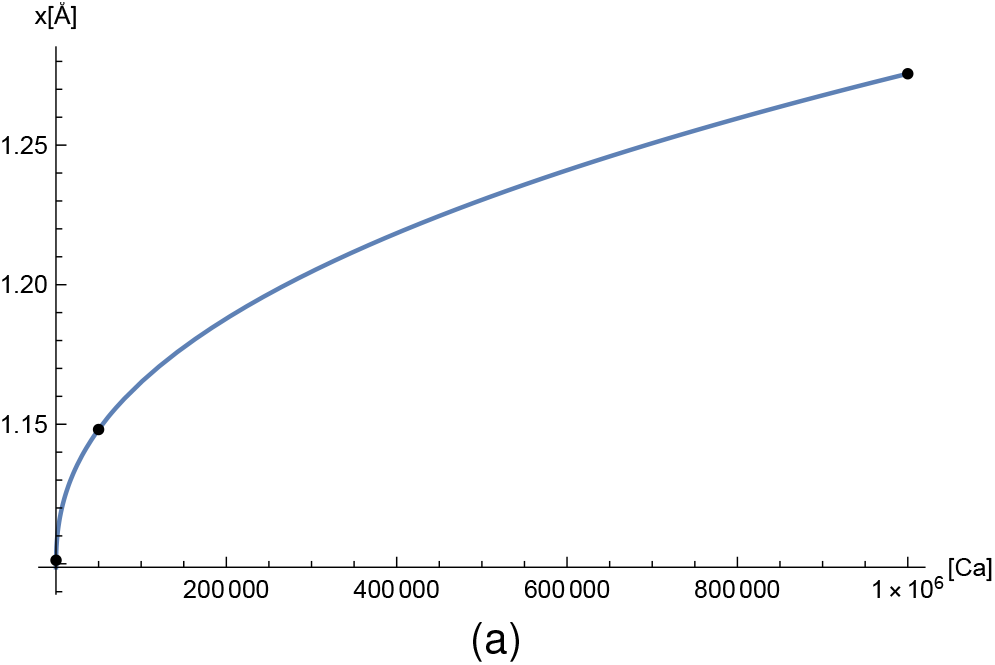
Impact of calcium concentration on the condensed layer thickness *x*_*c*_ − 1. From left to right, the black dots represent the points (50*nM*, 1.101), (50*µM*, 1.148), and (1*mM*, 1.276). The solid line represents the interpolating curve.

The values for the dielectric constant, layer thickness coming from the fraction of condensed ions, filament radius and surface charge density, as well as, the ion mobilities and bulk concentrations allow for calculations of the equivalent resistance values in both the longitudinal (axial) and transversal (radial) directions, the impedance (Table 2a), and conductivity in the axial direction (Table 3). Similarly, the ability to accumulate electric energy at different calcium concentrations is revealed in Table 2b along with the nonlinear parameter *b*^*eq*^.

**Table 2:** Impact of calcium concentration on (a) the equivalent transversal resistance 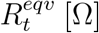, longitudinal resistance 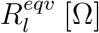, and impedance *Z* [Ω], (b) the free parameter *b*^*eqv*^ [1*/V*], and capacitance 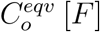.

| $[Ca^{2+}]$ | $R_t^{eq}[\times 10^7]$ | $R_l^{eq}[\times 10^8]$ | $Z[\times 10^9]$ |
| --- | --- | --- | --- |
| 50 nM | 1.322 | 1.836 | 4.82 |
| 50 $\mu M$ | 1.285 | 1.840 | 4.83 |
| 1 mM | 1.172 | 1.848 | 4.85 |

(a)
| $[Ca^{2+}]$ | $b^{eq}b$ | $C_o^{eq}[\times 10^{-17}]$ |
| --- | --- | --- |
| 50 nM | -0.178 | 1.801 |
| 50 $\mu M$ | -0.187 | 1.760 |
| 1 mM | -0.300 | 1.660 |

**Table 3:** Impact of calcium concentration on (a) the conductivity 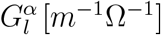 and effective cross section surface area 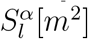 for the condensed layer (*α* = *CL*), and and (b) diffuse layer (*α* = *DL*), respectively. (c) Impact of calcium concentration on the equivalent conductance and effective cross section surface area 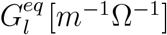 and 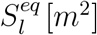 across the entire TLM.

| $[Ca^{2+}]$ | $G_l^{CL}$ | $S_l^{CL}[\times 10^{-18}]$ |
| --- | --- | --- |
| 50 *nM* | 0.14102 | 1.68702 |
| 50 $\mu M$ | 0.152744 | 1.76045 |
| 1 *mM* | 0.192547 | 1.96102 |

(a)
| $[Ca^{2+}]$ | $G_l^{DL}$ | $S_l^{DL}[\times 10^{-17}]$ |
| --- | --- | --- |
| 50 *nM* | 2.27769 | 1.28115 |
| 50 $\mu M$ | 2.26618 | 1.28326 |
| 1 *mM* | 2.23827 | 1.28899 |

(b)
| $[Ca^{2+}]$ | $G_l^{eq}[m^{-1}\Omega^{-1}]$ | $S_l^{eq}[\times 10^{-17}m^2]$ |
| --- | --- | --- |
| 50 *nM* | 0.132798 | $1.44985 \times 10^{-17}$ |
| 50 $\mu M$ | 0.143099 | $1.4593 \times 10^{-17}$ |
| 1 *mM* | 0.177295 | $1.48509 \times 10^{-17}$ |

### 3.2 Filament Characterization

Using the values for the equivalent resistances, capacitance, and impedance presented in the previous subsection, we characterized the dissipative parameters *µ*_2_ and *µ*_3_, along with the soliton amplitude Ω_*o*_ for voltage inputs of 0.05*V* and 0.15*V*. These values are tabulated in Table 4.

**Table 4:** Impact of input voltage *V*_*o*_ on the parameters *µ*_2_, *µ*_3_ and Ω_*o*_ for (a) 50 *nM* calcium concentration, and (b) 1*mM* calcium concentration.

| $V_o$ | $\mu_2$ | $\mu_3$ | $\Omega_o$ |
| --- | --- | --- | --- |
| 0.05 *V* | 0.0164547 | 0.913589 | 0.188774 |
| 0.15 *V* | 0.0164887 | 0.915474 | 0.326966 |

(a)
| $V_o$ | $\mu_2$ | $\mu_3$ | $\Omega_o$ |
| --- | --- | --- | --- |
| 0.05 *V* | 0.0144987 | 0.914032 | 0.24502 |
| 0.15 *V* | 0.0145507 | 0.917313 | 0.447798 |

To analyze the impact of the strong polarization on the water molecules surrounding the filament, we compared the propagation of ionic waves using two values for the dielectric constant in the condensed layer (Figure 5), namely *ϵ*_*CL*_ = 5 and *ϵ*_*CL*_ = 78.358. The latter is the one omitting the strong polarization of the electric dipole of the water molecules. The soliton profile snapshots for a voltage input of 0.15*V* with resting ([*Ca*^2+^] = 50 *nM*) and excited ([*Ca*^2+^] = 1 *mM*) neuron calcium concentrations are given in Figures 5a and 5b, respectively. Additionally, we compared the shape and attenuation of the ionic waves for the resting and excited neuron calcium concentrations to reveal the impact of the calcium concentration at a voltage input of *V*_*o*_ = 0.05*V*. The soliton profile snapshots corresponding to the dielectric constant *ϵ*_*CL*_ = 5 and *ϵ*_*CL*_ = 78.358 are presented in Figures 6a and 6b, respectively.

**Fig. 5:**
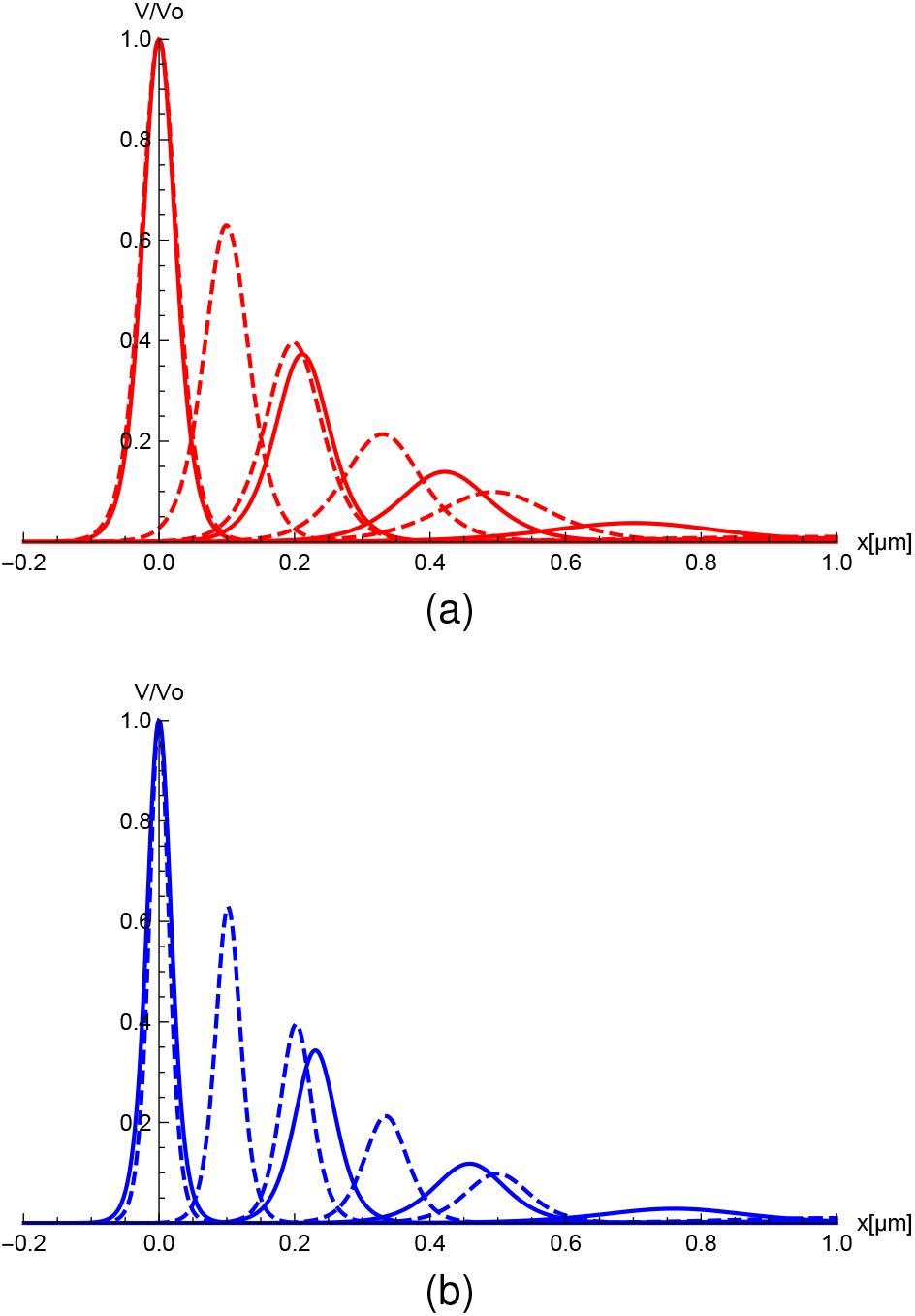
Impact of SL dielectric permittivity *ϵ*_*CL*_ on the soliton propagation for (a) 0.15*V* voltage input and 50*nM* calcium concentration and (b) 0.15*V* voltage input and 1 *mM* calcium concentrations. The solid and dashed red curves are for *ϵ*_*CL*_ = 5 and *ϵ*_*CL*_ = 78.358 SL dielectric permittivity, respectively. Soliton profile snapshots were taken at 0 *µs*, 0.85 *µs*, 1.71 *µs*, 2.85 *µs*, 4.27, *µs* and 8.54 *µs* for soliton peaks 1-6, respectively.

**Fig. 6:**
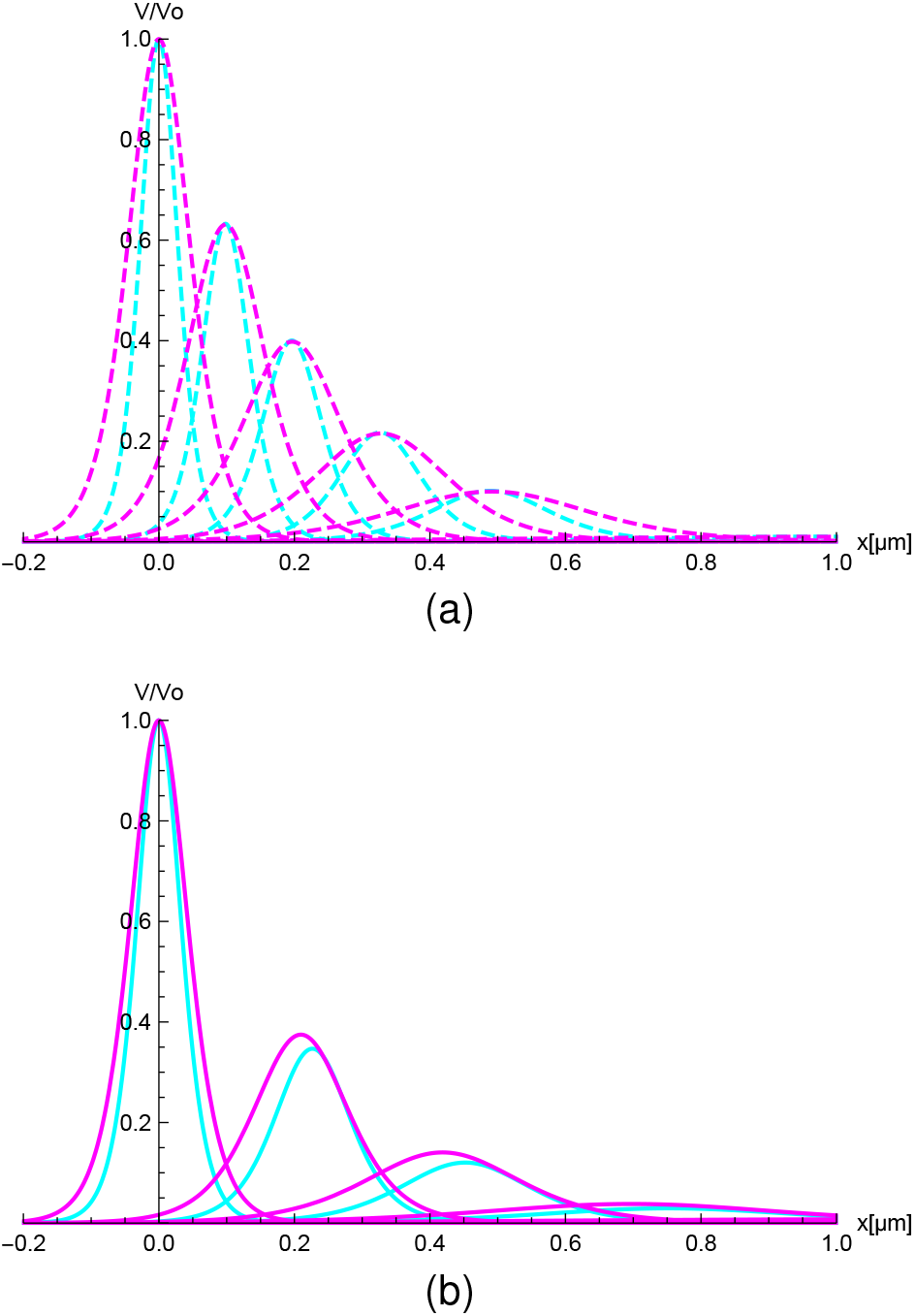
Impact of calcium concentration on the soliton propagation for (a) 0.15*V* voltage input and *ϵ*_*CL*_ = 78.358 SL dielectric permittivity (b) 0.15*V* voltage input and *ϵ* _*CL*_ = 5 SL dielectric permittivity. The magenta and cyan colors represent the soliton propagation for 50 *nM* and 1 *mM* calcium concentrations, respectively. Soliton profile snapshots were taken at 0 *µs*, 0.85 *µs*, 1.71 *µs*, 2.85 *µs*, 4.27, *µs* and 8.54*µs* for soliton peaks 1-6, respectively.

Using the proposed (more realistic) value for the stagnant layer dielectric constant *ϵ*_*CL*_ = 5, we analyzed the role of the input voltage and calcium concentration on the shape and attenuation of the ionic waves. In Figure 7a, we presented the soliton profile snapshots at the excited neuron calcium concentration for *V*_*o*_ = 0.05*V* and *V*_*o*_ = 0.15*V* voltage inputs, whereas in Figure 7b, we displayed the soliton profile snapshots at *V*_*o*_ = 0.15*V* voltage input for the resting and excited neuron calcium concentrations. Additionally, we analyzed the role of the input voltage and calcium concentration on the propagation of the ionic waves. In Figure 8, we present the velocity profiles as a function of time for the resting and excited neuron calcium concentrations and *V*_*o*_ = 0.05*V* and *V*_*o*_ = 0.15*V* voltage inputs. On the other hand, the velocity profile as a function of the calcium concentration for a voltage input of 0.15*V* is illustrated in Figure 9. Here we show both a double layer theory (Figure 9a) and triple layer theory (Figure 9b) for comparison purpose. Another important characterization of the filament comes from the vanishing time and maximum traveling distance of the soliton. The corresponding values for the resting and excited neuron calcium concentrations and *V*_*o*_ = 0.05*V* and *V*_*o*_ = 0.15*V* voltage inputs are tabulated in Table 5. A 3D image representation of a traveling soliton along an actin filament for a calcium concentration of 50 *nM* and voltage inputs of 0.15 *V* and 0.15 *V* is given in figures 10a and 10b to visualize the propagation velocity, shape and attenuation with both position and time. Figure 11 shows the soliton with a larger calcium concentration of 5 *mM* and a voltage input of 0.15 *V*.

**Table 5:** Impact of calcium concentration and input voltage on (a) soliton vanishing time and (b) maximum travel distance.

| $[Ca^{2+}]$ | $V_o = 0.05V$ | $V_o = 0.15V$ |
| --- | --- | --- |
| $50\text{ nM}$ | $7.89\text{ }\mu\text{sec}$ | $7.84\text{ }\mu\text{sec}$ |
| $1\text{ mM}$ | $7.30\text{ }\mu\text{sec}$ | $7.24\text{ }\mu\text{sec}$ |

(a)
| $V_o = 0.05V$ | $V_o = 0.15V$ |
| --- | --- |
| $0.969\text{ }\mu\text{m}$ | $0.968\text{ }\mu\text{m}$ |
| $0.967\text{ }\mu\text{m}$ | $0.965\text{ }\mu\text{m}$ |

**Fig. 7:**
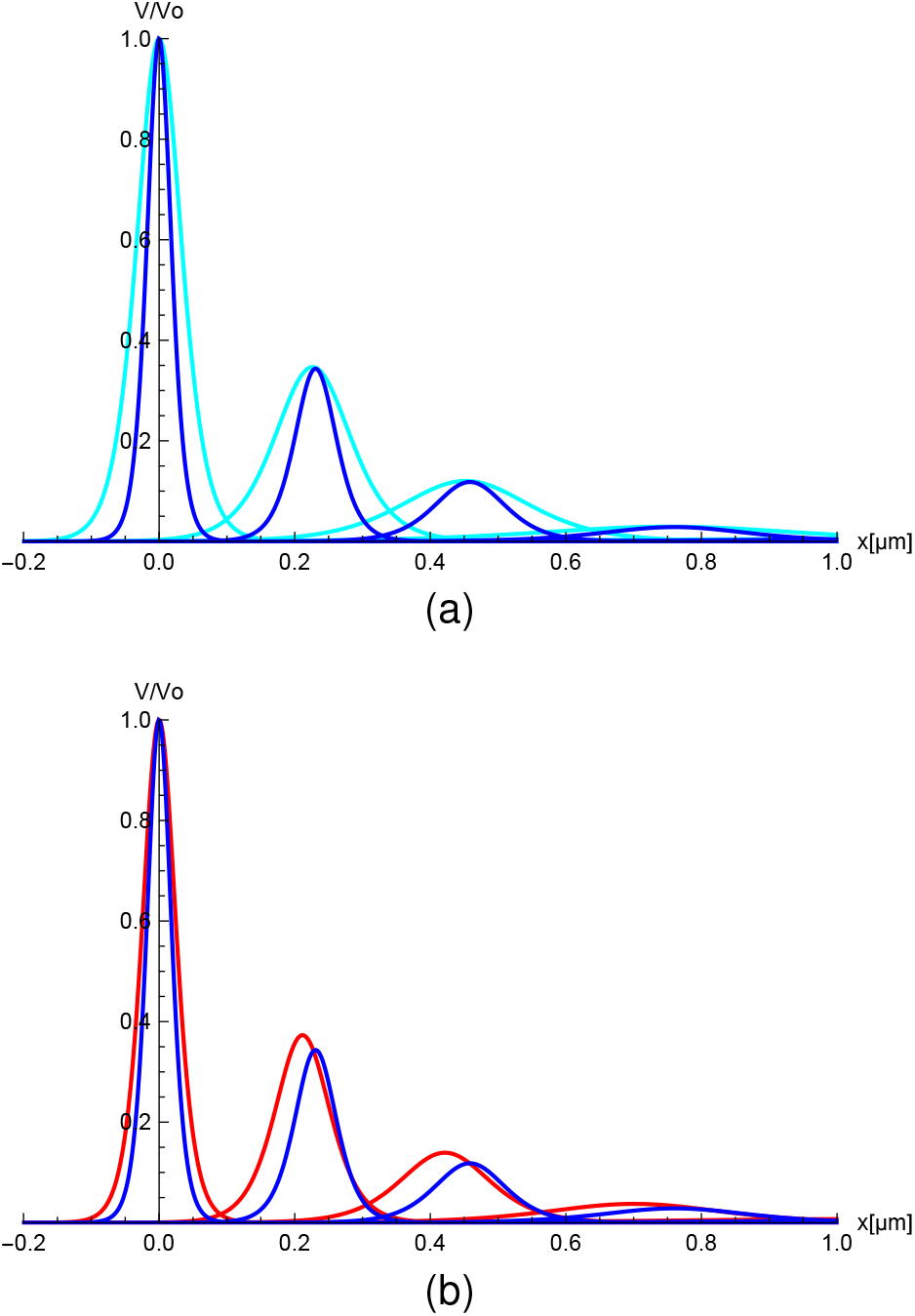
(a) Impact of input voltages on the soliton propagation for a calcium concentration of 1 *mM*. The cyan and blue colors represent the soliton propagation for 0.05*V* and 0.15*V* voltage inputs, respectively. (b) Impact of calcium concentration on the soliton propagation for 0.15*V* voltage input. Colors red and blue represent the soliton propagation for 50 *nM* and 1 *mM* calcium concentrations, respectively. Soliton profile snapshots were taken at 0 *µs*, 0.85 *µs*, 1.71 *µs*, 2.85 *µs*, 4.27, *µs* and 8.54*µs* for soliton peaks 1-6, respectively.

**Fig. 8:**
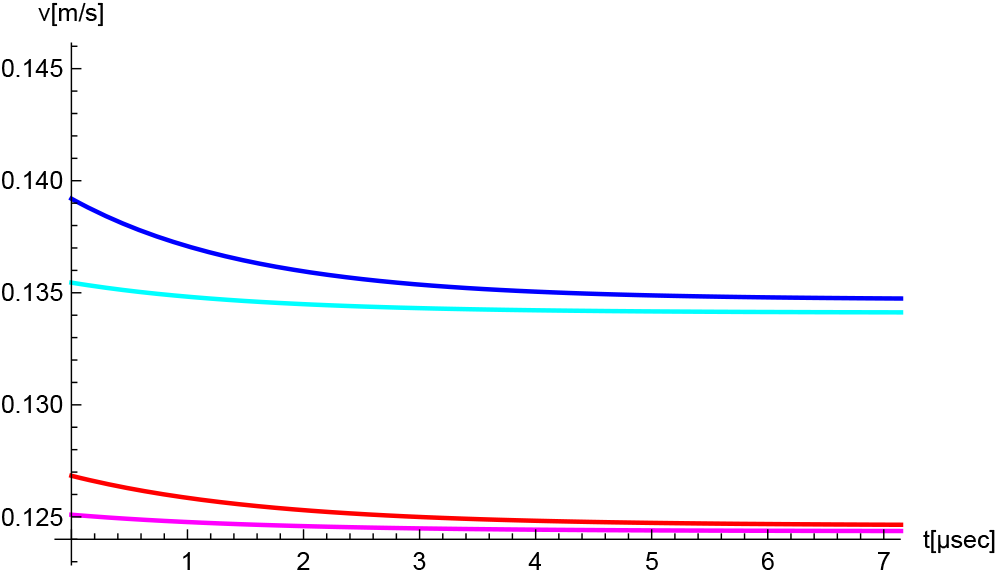
Impact of input voltage and calcium concentration on the soliton velocity profile. The cyan/magenta and blue/red lines represent the velocity profile for 0.05*V* and 0.15*V* voltage inputs, respectively.. Additionally, the red/magenta and blue/cyan lines represent the velocity profile for 50 *nM* and 1 *mM* calcium concentrations, respectively.

**Fig. 9:**
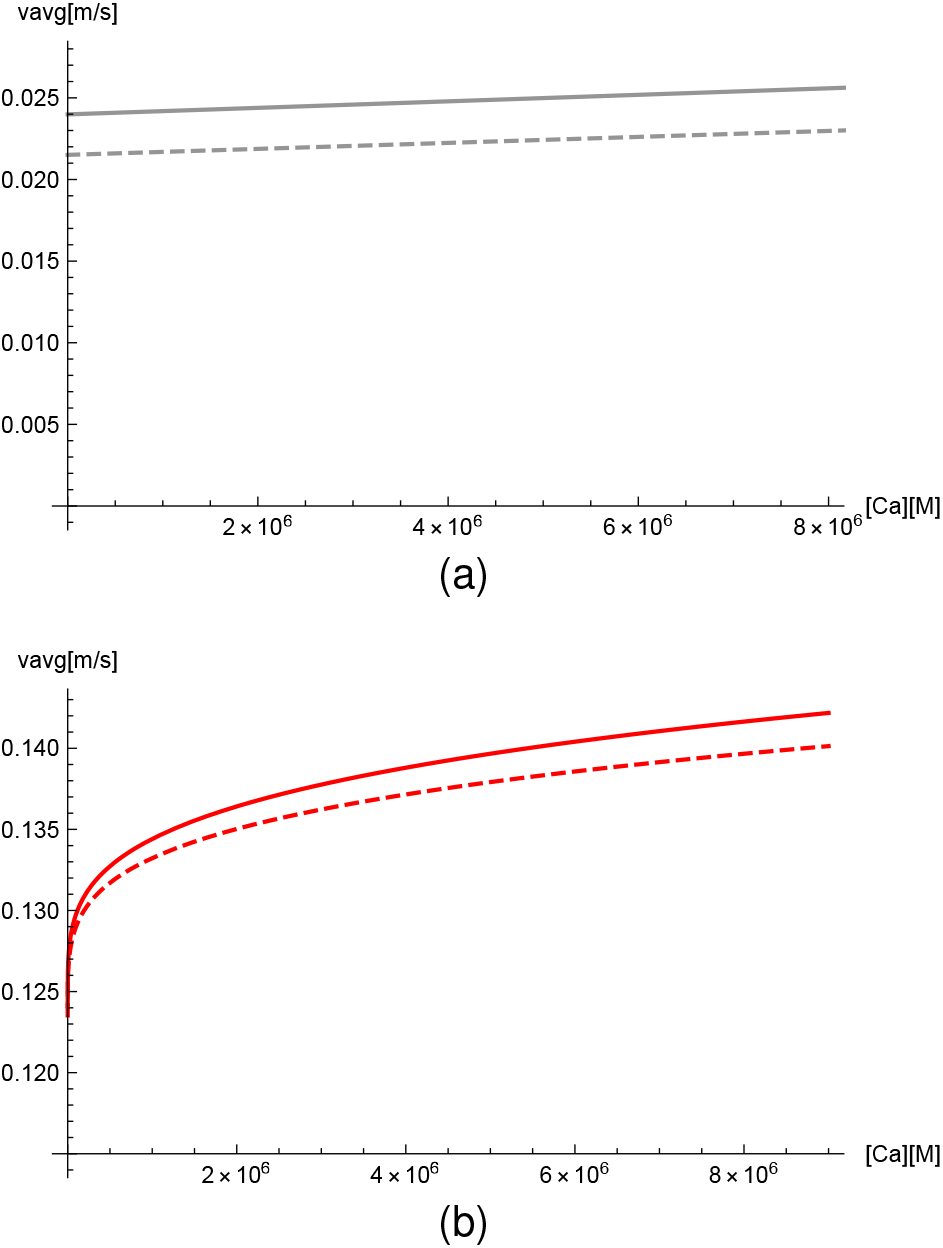
Average soliton velocity as a function of calcium concentration in (a) the double layer and (b) the triple layer models. Dashed and solid lines represent *V*_0_ = 0.05 *V* and *V*_0_ = 0.15 *V* voltage inputs, respectively.

**Fig. 10:**
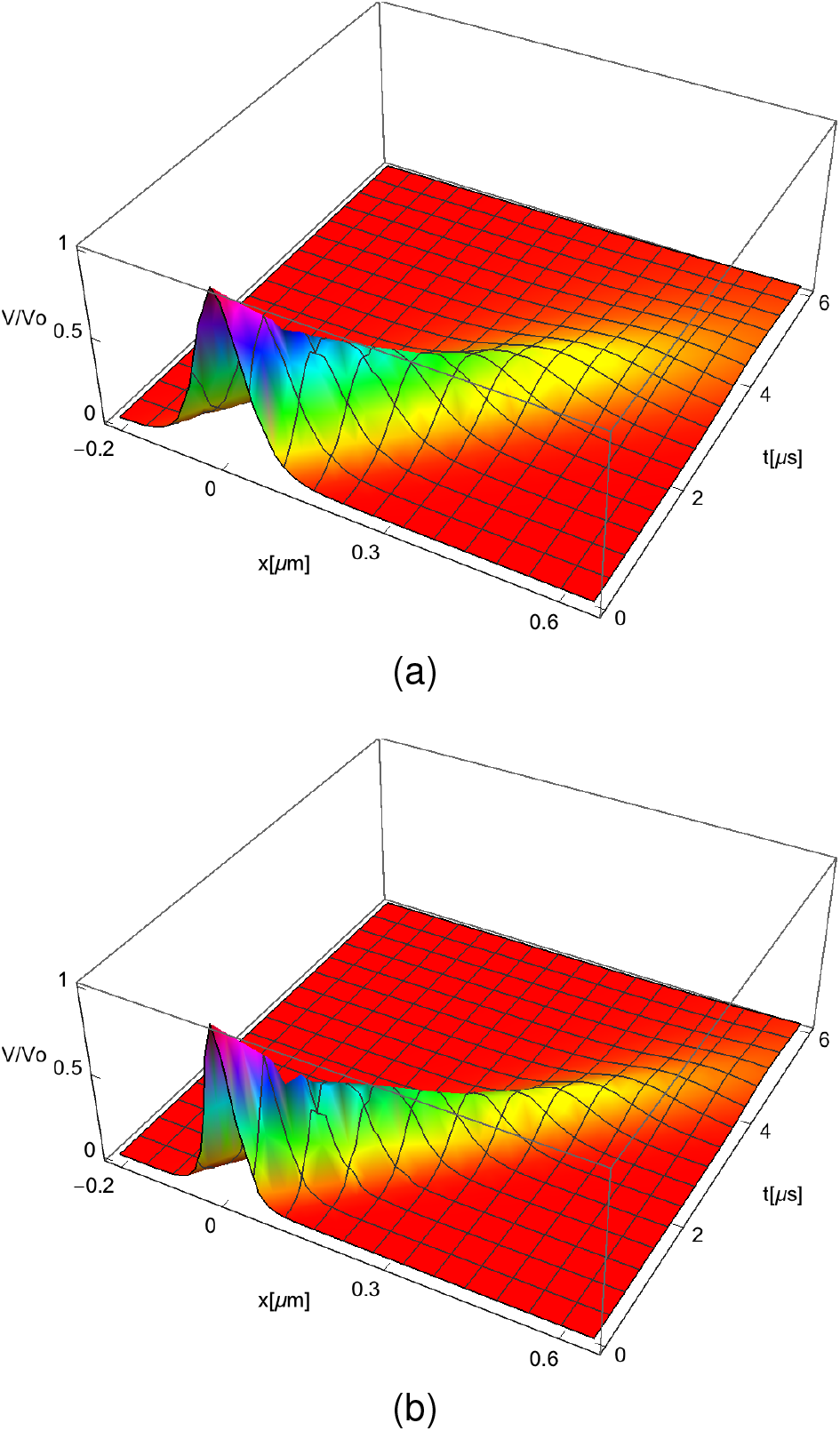
Normalized 3D soliton as a function of both position and time for 50*nM* calcium concentration and (a) *V*_0_ = 0.05*V* voltage input, (b) *V*_0_ = 0.15*V* voltage input.

**Fig. 11:**
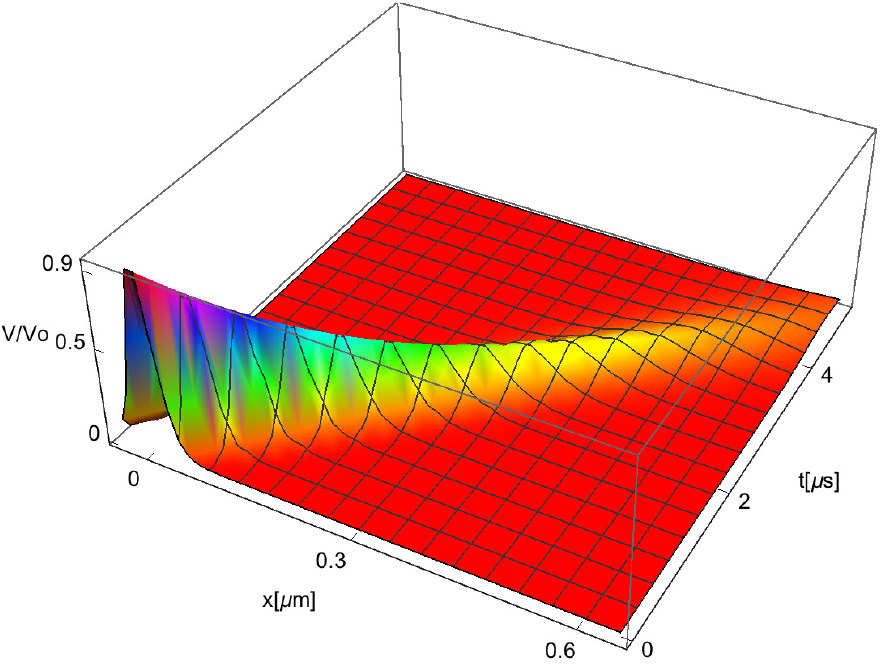
Normalized 3D soliton as a function of both position and time for 0.15 *V* input voltage and 1 *mM* calcium concentration.

### 3.3 Solitons Propagation during Neuron States Transition

The process of a conducting actin filament in a neuron going from the resting state to a fully excited state is revealed in Figure 12. The pink curve represents a neuron at a resting calcium concentration of 50 *nM* after a stimulus of 0.05*V* was applied. The blue curve shows how a packet of ions travels down the filament at the fully excited state (∼ 0.15*V*) with a calcium concentration of 1 *mM*.

**Fig. 12:**
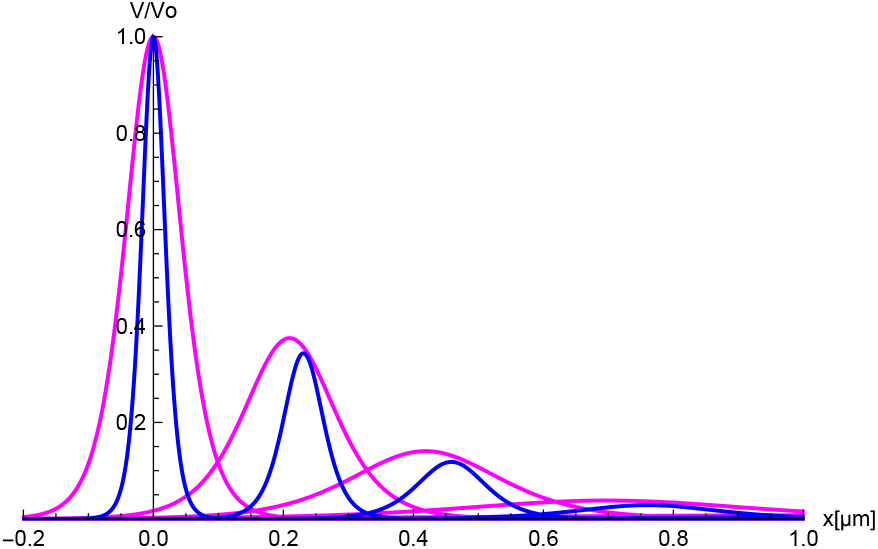
Solition propagation in the process of a true resting state to a fully excited neuron. The pink curve represents 0.05 *V* at 50 *nM* calcium concentration. The blue curve is for 0.15 *V* at 1 *mM* calcium concentration. Soliton profile snapshots were taken at 0 *µs*, 0.85 *µs*, 1.71 *µs*, 2.85 *µs*, 4.27, *µs* and 8.54 *µs* for soliton peaks 1-6, respectively.

## 4 Discussion

The triple layer model reveals a widening of the condensed layer for larger bulk concentrations of calcium (see Figure 4) since more ions are attracted to the negatively charged actin filament surface. In particular, when a resting neuron is excited, the concentration of condensed ions increases 0.1018 and consequently, the stagnant layer thickness enlarges 14% (Table 1a). The comparison of our results with the (conventional) double layer Manning’s theory shows a decrease in one order of magnitude lower in the values obtained for the fraction of condensed ions. Additionally, Manning’s theory predicts the same fraction of condensed ions for the resting and excited neuron states (Table 1b). These discrepancies arises from the different dielectric constant value used near the filament surface, as well as, the accuracy in the approximations used in the NLPB theory to calculate the electric potential. The drawback in using an overestimated (bulk) value for the dielectric constant used near charged colloidal systems in the double layer model was already pointed it out in previous work on condensed ion current along conducting surfaces (Dhont and Kang, (2011)). In contrary, we consider the strong polarization of the electric dipole of the water molecules in the stagnant layer arising from the local electric field generated by the filament surface charge density. As a result, the value of the dielectric constant in the stagnant layer is one order of magnitude lower than that used in the double layer model. Our approach (*ϵ* _*CL*_ = 5 in the stagnant layer and *ϵ* _*DL*_ = 78.358 in the diffuse layer) is in agreement with the (more realistic) sigmoid-like distance-dependent dielectric constant predicted by more sophisticated Polar classical DFT-like theories (Warshavsky and Marucho, (2016)). Additionally, we use the NLDH approximation which has been shown to be more accurate than the DH approximation used by Manning theory (Lamm and Pack, 2010). Unlike the DH approximation, the NLDH approximation is able to predict mono- and multivalent ion condensation, and it captures the dependence of the fraction of condensed ions on the bulk concentration. Whereas, a non dependent ion bulk concentration for multi-valent ion condensation is predicted by Manning’ theory when using an ad-hoc free-bound two ion states assumption along with the DH approximation.

The triple layer model proposed in this work provided not only a more accurate description of the electric potential and ionic distribution functions near the filament surface but also a more realistic characterization on the monomeric conductivity and capacitance properties. It is worth noting that the triple layer electric circuit can be reduced to a double layer electric circuit with effective (equivalent) elements using the parallel and series connection properties of the SL and DL resistances and capacitances (Table 2). Our results show that the monomeric transversal (radial) and longitudinal (axial) resistances, as well as, impedance and bare capacitance decrease with increasing calcium bulk concentrations. Whereas, it has the opposite behavior on the non linear capacitance parameter. This means that G-actin proteins in the excited neuron state display higher non-linear accumulation of charge and conductivity than in the resting state. The role of the condensed ions in the stagnant layer can be further elucidated by analyzing its contribution to the effective monomeric axial conductivity at different calcium bulk concentrations (Table 3). Our results show a substantial increase of the SL conductivity and cross-section area with increasing calcium bulk concentrations since a higher fraction of ions are condensed and the thickness of the stagnant layer is enlarged. Whereas, a slight increase in the conductivity and decrease of the cross-section area is seen for the DL (Table 3a). Overall, the effective monomeric conductivity is dominated by the contribution coming from the DL since its conductivity and cross-section area are one order of magnitude higher than those values corresponding to the SL (Table 3c).

We also investigated the role of the SL dielectric constant in the soliton propagation profiles. At the resting neuron calcium concentration, solitons travel faster, display higher attenuation, as well as, broader distribution around the peaks when they propagate in a low dielectric medium *ϵ* _*CL*_ = 5 (see solid versus dashed curves in Figure 5a). This effect is even more pronounced for neurons in the excited state (see solid versus dashed curves in Figure 5b). Additionally, solitons in a low dielectric medium travel faster, display higher attenuation, as well as, broaden distribution around the peaks for neurons in the excited state (see Figure 6b). A different, probably incorrect, scenario is visualized on the soliton propagation profiles when an overestimated (bulk) value is used for the SL dielectric constant. Solitons propagating in a high dielectric medium travel at the same velocity (peak positions) and with the same attenuation (height of the peaks) for both the resting and excited neuron calcium concentrations. Meanwhile, solitons in the excited neuron state display narrower distributions around the peaks (see Figure 6a).

In this work, we considered low-range electrical stimulus to simulate voltage clamp experiments. Our results show broaden soliton distributions around the peaks and neglectable changes on the attenuation with increased voltage inputs (see Figure 7a). Furthermore, the information obtained on the time-dependent velocity profiles revealed a soliton deceleration, which is more pronounced in the excited neuron state (see figure 8).

Our analysis on the impact of calcium bulk concentrations on the soliton propagation profiles show that solitons travel faster, display higher attenuation, as well as, broader distributions around the peaks in neurons in the excited state (see figure 7b). Furthermore, the time average soliton velocity logarithmically increases with increasing calcium bulk concentrations (Figure 9b). This behavior is due to the stagnant layer since it was not observed in the previous results using a double layer model (see figure 9a). The stagnant layer also generates an increase in one order of magnitude in the average soliton velocity.

In general, our results predict solitons traveling an average maximum distance around 1.0 *µm* along the actin filament for both the excited and resting neuron states. Whereas, the soliton travels around 1 *µs* longer in the excited neuron state (see Table 5a).

Further analysis was recently carried out at higher voltage inputs showing significant changes on the average soliton velocity, attenuation and maximum traveling distance at different calcium bulk concentrations for voltage inputs larger than 0.5V (results not shown in results section).

## 5 Conclusions

In this article, we introduced a novel electrical triple layer model and used a non-linear Debye-Huckel theory in a non linear, dissipative, electrical transmission line to elucidate the role of the ion condensation and the high polarization of the immobile water molecules in the propagation of ionic calcium waves along F-actin. As a unique feature, the multi-scale approach accounts for the physicochemical properties of each monomer in the filament in terms of an electric circuit model containing monomeric flow resistances and ionic capacitances in both the condensed and diffuse layers. In our studies we characterized the biocylindrical actin filament model using a high resolution molecular structure. We considered the resting and excited state of a neuron using 0.1 *M KCl* + 50 *nM CaCl*_2_ and 0.1 *M KCl* + 1 *mM CaCl*_2_ electrolyte mixtures, respectively. Additionally, we used 0.05*V* and 0.15*V* voltage inputs to investigate ionic waves in voltage clamp experiments on actin filaments.

Our results showed that G-actin proteins in the excited neuron state displayed higher nonlinear accumulation of charge and conductivity than in the resting state. This is due in part to a larger fraction of condensed calcium ions and increase in width of the condensed layer thickness when increasing the calcium concentrations. Additionally, our results revealed a less disperse wave packet of calcium ions, as well as, a faster propagating soliton for larger calcium concentrations and input voltages. Consequently, there are more calcium ions transmitted reaching the opposite end of the actin filament faster.

We conclude that the different conduction properties arising from the condensed and diffuse layers in the ETL model provide an avenue of delivery through cytoskeleton filaments for larger charged cationic species. This new direct path for fast divalent ion transport inside neurons might be of fundamental importance for neurological diseases that connect different compartments of the neuron to the soma. It might be also relevant for the development of reliable, highly functioning small devices with biotechnological applications such as bionanosensors and computing bionanoprocessors (Patolsky, Weizmann, and Willner, 2004; Zhou et al., 2010; Agarwal and Hess, 2010).

## Acknowledgments

This work was supported by NIH Grant 1SC1GM127187-01.

## Availability of data and code

Codes used for this study can be requested from the corresponding author.

## Ethics approval

Not Applicable

## Consent to participate

Not Applicable

## Consent for publication

Not Applicable

## Appendix

### Validation on the mean electrostatic potential approximation

In Figures 13a and 13b we display in green color the approximate mean electrostatic potential solution used in the diffuse layer for [*Ca*] = 50*nM* and [*Ca*] = 1*mM* calcium concentrations, respectively. For comparison purposes, we also include the orange and blue lines which correspond to the exact nonlinear Debye-Huckel and the linear Debye-Huckel solutions, respectively.

**Fig. 13:**
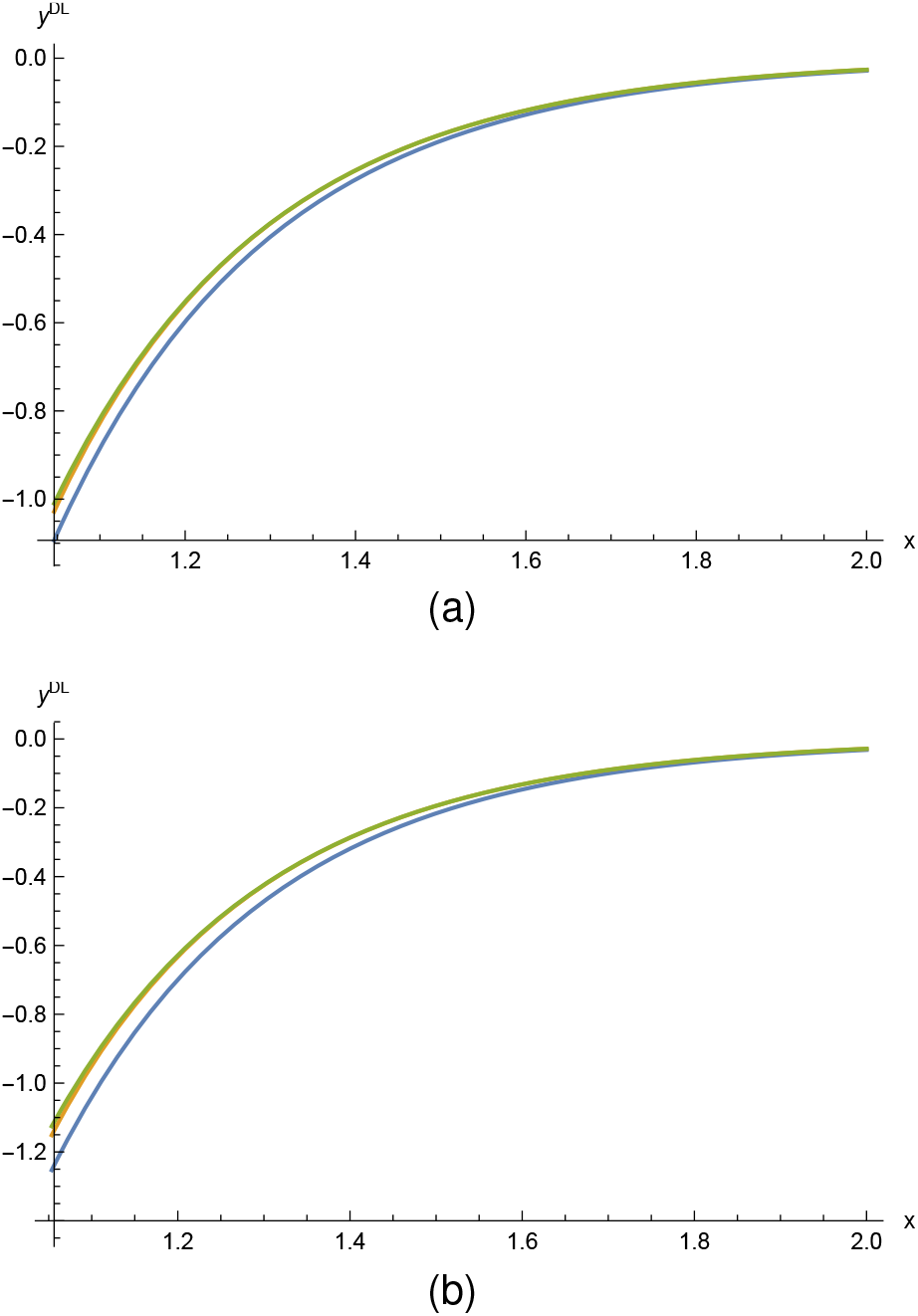
Figure (a) is for [*Ca*] = 50*nM* and figure (b) is for calcium concentration [*Ca*] = 1*mM*. The orange, green, and blue lines correspond to the exact nonlinear Debye-Huckel, the nonlinear Debye-Huckel approximation (this work), and the linear Debye-Huckel solutions.

### Non trivial contributions to the longitudinal conductivity coming the the diffuse layer

The explicit expression for 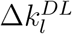 reads

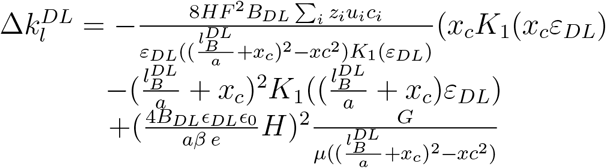

where

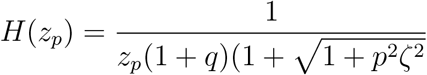

and

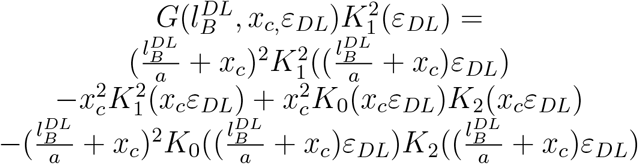

#### Condensed layer capacitance parameters

The expressions for the parameters *a*_0_, *a*_1_, *a*_2_, and 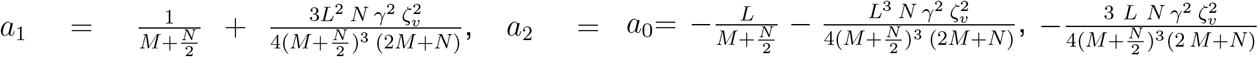, and 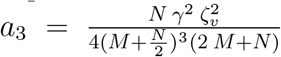, where 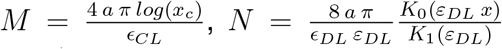 and 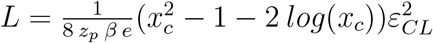.

#### Impedance expression for the triple layer model

The expression for the impedance in the triple layer model reads

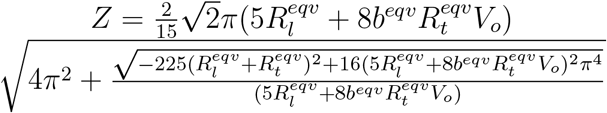

For *R*_*t*_ *≪ R*_*l*_, this expression recovers the approximation *Z* ≃ 25.1128*R*_*l*_ used in our previous work on the EDL in which the transversal resistance was neglectable.

